# Candidate stress biomarkers for queen failure diagnostics

**DOI:** 10.1101/2020.02.27.961847

**Authors:** Alison McAfee, Joseph Milone, Abigail Chapman, Leonard J Foster, Jeffery S Pettis, David R Tarpy

## Abstract

**Background:** Queen failure is a persistent problem in beekeeping operations, but in the absence of overt symptoms it is often difficult, if not impossible, to ascertain the root cause. Stressors like heat-shock, cold-shock, and sublethal pesticide exposure can reduce stored sperm viability and lead to cryptic queen failure. Previously, we suggested candidate protein markers indicating heat-shock in queens, which we investigate further here, and tested new stressors to identify additional candidate protein markers.

**Results:** We found that heat-shocking queens for upwards of one hour at 40 °C was necessary to induce significant changes in the two strongest candidate heat-shock markers, and that relative humidity significantly influenced the degree of activation. In blind heat-shock experiments, we tested the efficiency of these markers at assigning queens to their respective treatment groups and found that one marker was sufficient to correctly assign queens 75% of the time. Finally, we compared cold-shocked queens at 4 °C and pesticide-exposed queens to controls to identify candidate markers for these additional stressors, and compared relative abundances of all markers to queens designated as ‘healthy’ and ‘failing’ by beekeepers.

**Conclusions:** This work offers some of the first steps towards developing molecular diagnostic tools to aid in determining cryptic causes of queen failure. Further work will be necessary to determine how long after the stress event a marker’s expression remains changed and how accurate these could be in the field.

## Introduction

According to surveys in Canada [1] and the US [2, 3], one of the most frequently reported causes of honey bee (*Apis mellifera*) colony failure is ‘poor queens.’ Unfortunately, the underlying factors leading to queen failure are usually difficult, if not impossible, to determine. For example, previous research has found that heat-shock, cold-shock, and pesticide exposure all decrease the viability of sperm stored in queen spermathecae [4, 5], which would theoretically lead to the same colony-level symptoms: inconsistent brood patterns, poor population build-up, and sometimes atypical drone-laying. If beekeepers cannot diagnose the root cause of queen failure, they are less equipped to identify the source of stress and are unable to make evidence-based management decisions to mitigate or eliminate the stress in the future.

In colonies, honey bees are normally adept at thermoregulating [6], but queens are vulnerable to heat-shock and cold-shock events during shipping [4, 7], where they are regularly transported long distances via ground or air cargo in poorly thermoregulated environments [8]. In a colony, worker bees can try to lessen the effects of overheating by collecting water and fanning to achieve evaporative cooling, or by heat-shielding, using their own bodies as heat-blocking insulation [6, 9]. Likewise, they can mitigate chilling by vibrating their wing muscles to generate heat [6]. However, in small queen cages used for shipping, there is often limited (if any) water, poor ventilation, and too few workers to effectively cool the queen. Likewise, if temperatures drop, workers can generate heat by vibrating their wing muscles; however, this strategy is not effective in small cages with few bees. We have previously included temperature loggers in long distance queen shipments and found that both hot and cold temperature spikes regularly occur [4, 7]. Some commercial queen suppliers include temperature loggers in their shipments as a quality control measure, but these are typically only large-order, international deliveries, and it is up to the discretion of the supplier. Worryingly, some evidence suggests that temperature spikes can even occur inside colonies during extreme heat-waves [7, 10] – conditions that are projected to increase in frequency, severity, and duration with climate change [11, 12].

Since queens are fed a strict diet of worker glandular secretions (*i.e*, they do not consume potentially contaminated flower products) [13-15], they are normally buffered from direct oral xenobiotic exposure [16, 17]. However, pesticides (including miticides, fungicides, herbicides, and insecticides) accumulate in wax [18, 19] and thus can pose a contact exposure hazard to queens. A cumulative hazard quotient (HQ) is one way to interpret risk during co-exposures to multiple pesticides simultaneously and employs a solely additive approach to incorporate the dose and respective toxicity of all pesticides in a mixture to estimate hazard [20]. In a survey of commercial colonies engaged in agricultural pollination, Traynor *et al*. found that the mean hazard quotient of the cumulative pesticide ‘exposome’ in wax was 2,155 [18]. Those colonies experiencing ‘queen events’ (*i.e*., the colony was queenless, had queen cells, or a virgin queen) had an average wax HQ of ∼3,500, while queenright colonies had average wax HQ of ∼1,700. While this is only a correlational observation, other researchers have found increased frequencies of queen events when colonies were given pollen patties with clothianidin [21] or a mix of clothianidin and thiamethoxam [22], both of which are neonicotinoid pesticides. Neonicotinoids can also reduce stored sperm viability [5], which is a possible mechanism leading to queen failure events.

Our ultimate goal is to identify queen stress biomarkers that can help distinguish different causes of queen failure in a single diagnostic test (**Figure 1**). As a first step towards that goal, here we heat-shocked, cold-shocked, and pesticide-stressed queens experimentally and compared protein expression profiles of their spermathecae to controls. The spermathecae were chosen because if a queen is sent for laboratory analysis, the spermathecae will likely be dissected to determine sperm viability and sperm count metrics, and we have previously shown that the remaining tissue sample is amenable to proteomics by mass spectrometry. Therefore, it is an economical and biologically relevant choice. We also surveyed protein expression profiles in 125 ‘imported,’ ‘healthy,’ and ‘failed’ queens to look for overlap between their expression patterns and the candidate biomarkers. Finally, we conducted a blind heat-shock trial to test the efficacy of using the top two candidate heat-shock biomarkers to assign queens to their relevant treatment groups. This work forms the foundation on which to further develop and refine these candidate queen stress biomarkers for eventual diagnostic use.

**Figure 1.**
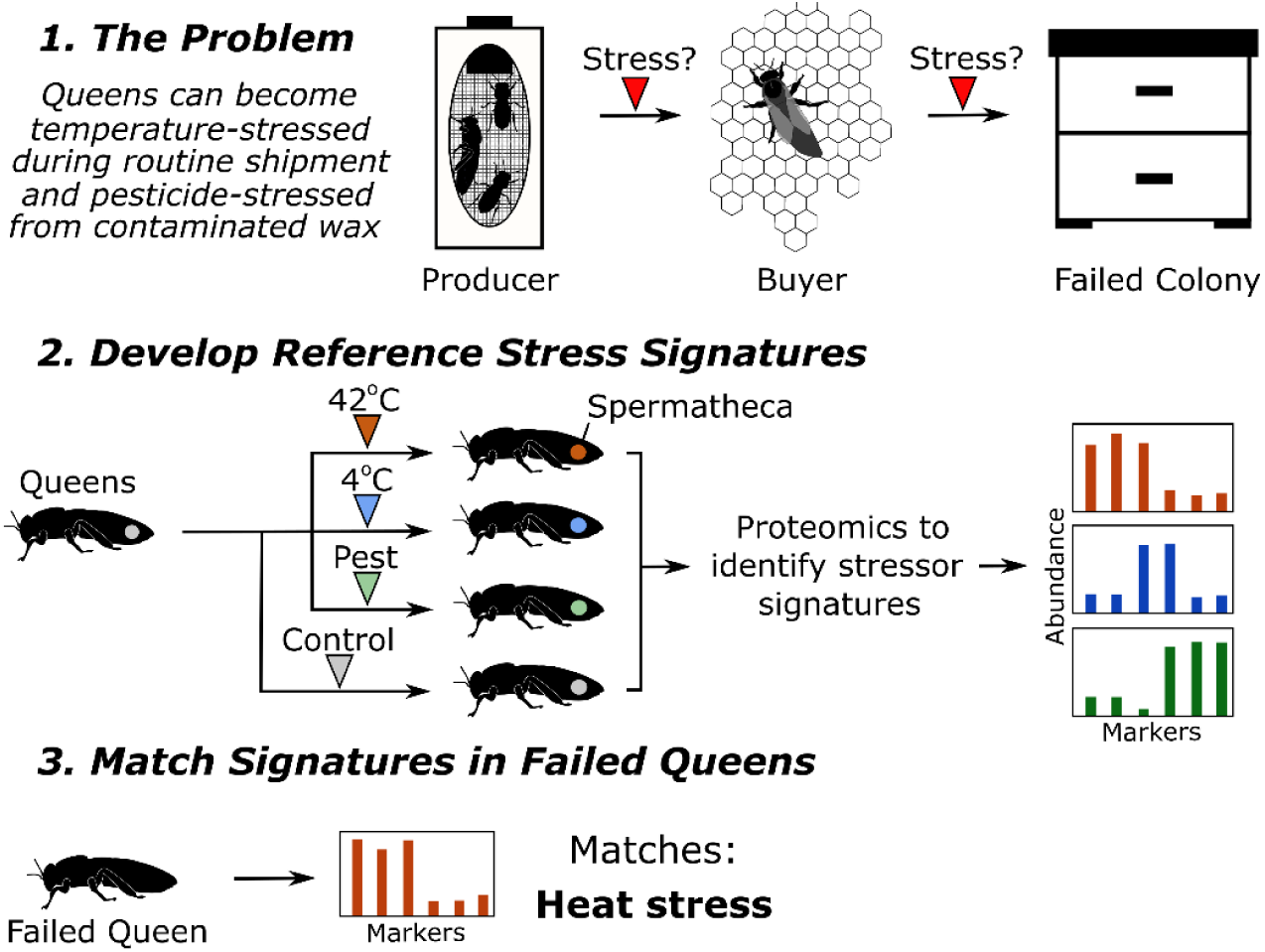
Schematic of experimental design for identifying queen stress biomarkers. Queen silhouettes adapted from McAfee *et al*. (2019), reuse permitted under CC BY 4.0.

## Results

### Temperature stress

Previously, we heat-shocked queens and used comparative proteomics to suggest candidate spermathecal protein markers [7]. To expand our stressor scope, here we also compared cold-shocked queens to controls and investigated the proposed heat-shock markers in more detail through an exposure time-course. Of the 2,094 quantified proteins, 21 were differentially expressed between cold-shocked (2 h at 4 °C) and control queens (n = 5 and 7, respectively), and only three of those were upregulated in the cold-shock group (**Figure 2a;** Student’s *t* test, 5% false discovery rate (FDR), Benjamini-Hochberg correction). GO term enrichment by the gene score resampling approach, which does not depend on p value thresholds [23], shows that acyl transferase activity was significantly upregulated with cold-shock, whereas oxidoreductase activity was significantly downregulated, among others (**Figure 2b**; 5% FDR, Benjamini-Hochberg correction). We anticipate that upregulated, rather than downregulated, proteins will generally be the most practical biomarkers because they are less likely to be affected by instrumental limits of detection and should not be as sensitive to false positives (*e.g*., from sample degradation). Therefore, we selected the top two most significantly upregulated proteins, XP_026296654.1 (leucine-rich repeat transmembrane neuronal protein 4-like) and XP_395122.1 (probable tRNA N6-adenosine threonylcarbamoyltransferase) as candidate cold-shock biomarkers.

**Figure 2.**
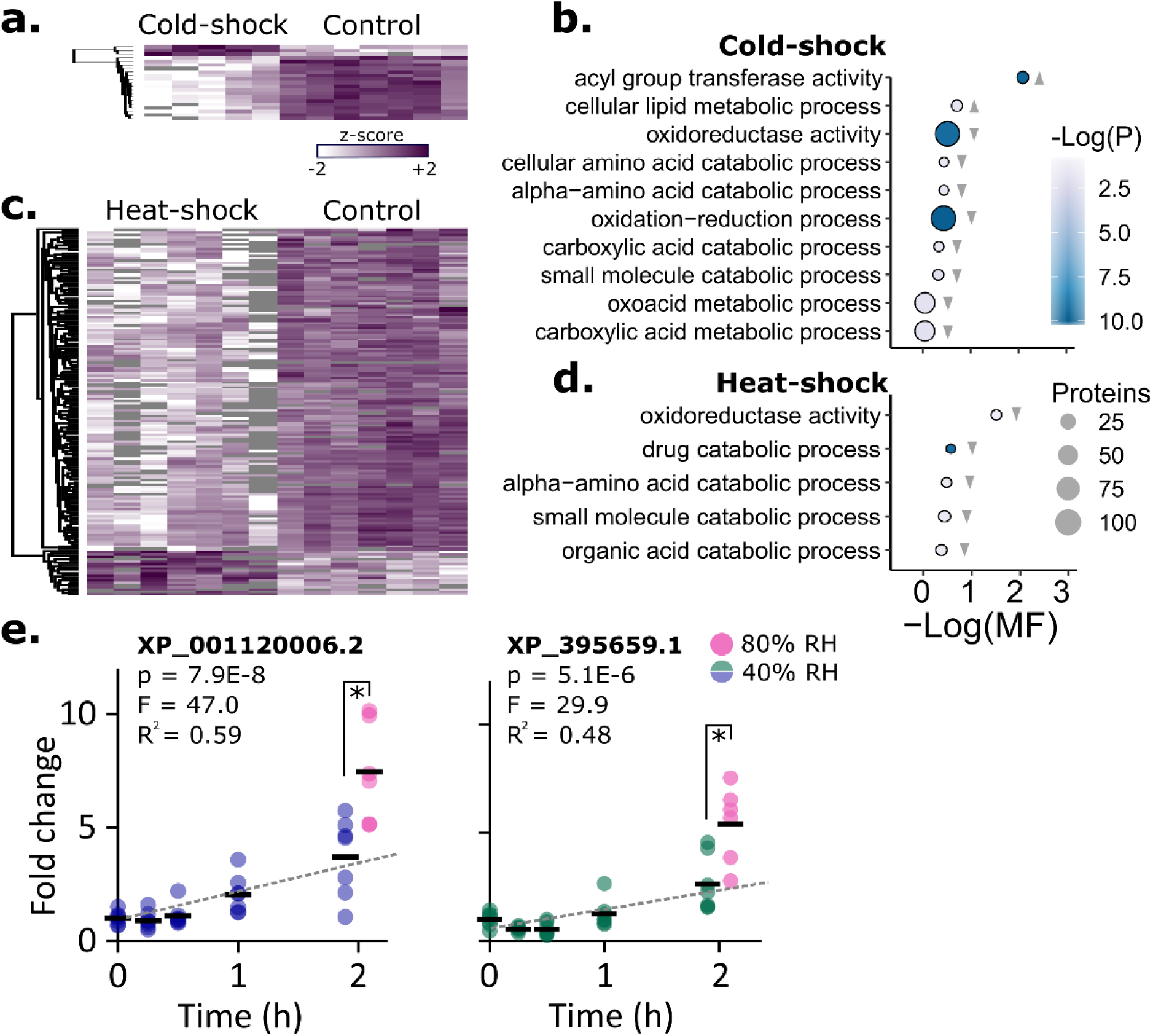
Effects of heat-shock and cold-shock on spermathecal protein abundance. **A)** Protein expression changes induced by cold-shock at 4 °C for two hours (only significant proteins surviving 5% FDR threshold by the Benjamini-Hochberg method are shown). The two most significantly upregulated proteins are XP_026296654.1 (leucine-rich repeat transmembrane neuronal protein) and XP_395122.1 (probable tRNA N6-adenosine threonylcarbamoyltransferase). Each column represents the spermatheca from one queen. **B)** GO term enrichment analysis. Grey triangles indicate the direction of expression driving the enrichment (relative to control). Only GO terms passing a 5% false discovery cut-off are shown (Benjamini-Hochberg method). MF stands for multifunctionality score (higher values are less multifunctional). Circle size is proportional to the number of proteins belonging to the GO term. **C)** Protein expression changes induced by heat-shock at 40 °C for two hours. **D)** Enrichment analysis. **E)** Temporal changes in expression of two previously identified candidate heat-shock biomarkers. Expression was significantly influenced by time and relative humidity (RH). Black bars indicate the mean.

The two most significant candidate heat-shock biomarkers we identified in our previous experiment [7] were the small heat-shock proteins XP_395659.1 and XP_001120006.2, and here we confirm that these two proteins are consistently upregulated across queens from independent origins. Both proteins were again globally differentially expressed after a 2 h, 40 °C exposure relative to negative controls (5% FDR, Benjamini-Hochberg, n = 7 each), and this response was significantly higher at 80% relative humidity (RH; n = 7) compared to 40% RH (**Figure 2e;** n = 6; XP_001120006.2: df = 11, p = 0.0055, T_stat_ = 3.44; XP_395659.1: df = 11, p = 0.0075, T_stat_ = 3.26). Holding the RH at a constant 40%, expression of XP_001120006.2 (linear regression, df = 34, p = 7.9×10^−8^, F = 47.0) and XP_395659.1 (df = 33, p = 5.1×10^−6^, F = 29.9) significantly depended on exposure duration, with clear differences not emerging until the 1 h time point.

### Pesticide stress

Temperature is one of many stress factor that can influence queen quality, as pesticide exposure can also reduce sperm viability and lead to queen failure [5, 21, 22]. To evaluate effects of direct contact pesticide exposure on spermathecal protein expression, we applied 2 µl of either acetone alone, acetone with 20 ppb imidacloprid (a neonicotinoid insecticide), a pesticide “cocktail” (which does not contain any neonicotinoids) diluted in acetone to achieve a hazard quotient of ∼511 (**Table 1**). This HQ is considerably lower than the mean wax HQ in commercial colonies (2,155), which was intentional because we expect the transfer of toxins from wax to the queen to be inefficient and we also sought to examine the queen biomarkers resulting from sublethal pesticide exposure. The imidacloprid dose is also an equivalent amount to the lowest dose Chaimanee *et al*. [5] tested on queens (2 ul of 20 ppb imidacloprid), which they found caused a significant drop in sperm viability. No queens died during these experiments.

**Table 1.** Concentrations and hazard quotients (HQ) for components of the pesticide cocktail

| Pesticide | Mode of action | Group | Median detection (ppb)* | Target ppb | Contributing HQ |
| --- | --- | --- | --- | --- | --- |
| Coumaphos | Acetylcholine esterase inhibitor | Acaricide | 943 | 314 | 53.00 |
| tau-Fluvalinate | Sodium channel modulator | Acaricide | 4310 | 1437 | 332.56 |
| 2,4-DMPF** | Octopamine receptor agonist | Acaricide (metabolite) | 304 | 101 | 1.35 |
| Chlorothalonil | Multisite activity | Fungicide | 361 | 120 | 1.08 |
| Chlorpyrifos | Acetylcholine esterase inhibitor | Insecticide | 2.7 | 1 | 11.81 |
| Fenpropathrin | Sodium channel modulator | Insecticide | 16.8 | 6 | 112 |
| Pendimethalin | Inhibition of microtubule assembly | Herbicide | 5.3 | 2 | 0.02 |
| Atrazine | Inhibition of photosynthesis | Herbicide | 5.4 | 2 | .01 |
| Azoxystrobin | Cytochrome bc1 inhibitor | Fungicide | 5.1 | 2 | .01 |
| **Total HQ** |  |  |  |  | **511** |

\* Median wax concentrations and LD50s as determined by Traynor *et al.* (2016) [18]
\*\* 2,4-Dimethylphenyl formamide (Amitraz degradate)

We found stark differences between cocktail-treated and untreated queens, with intermediate effects of acetone alone (**Figure 3a;** n = 7, 10, and 8, respectively). The effect of acetone in our solvent control group was significant (653 proteins were differentially expressed between acetone-treated and untreated queens at 5% FDR, Benjamini-Hochberg correction), indicating that, although commonly used, this organic carrier is far from benign. However, our interest is in proteins specifically altered by the pesticide treatments.

**Figure 3.**
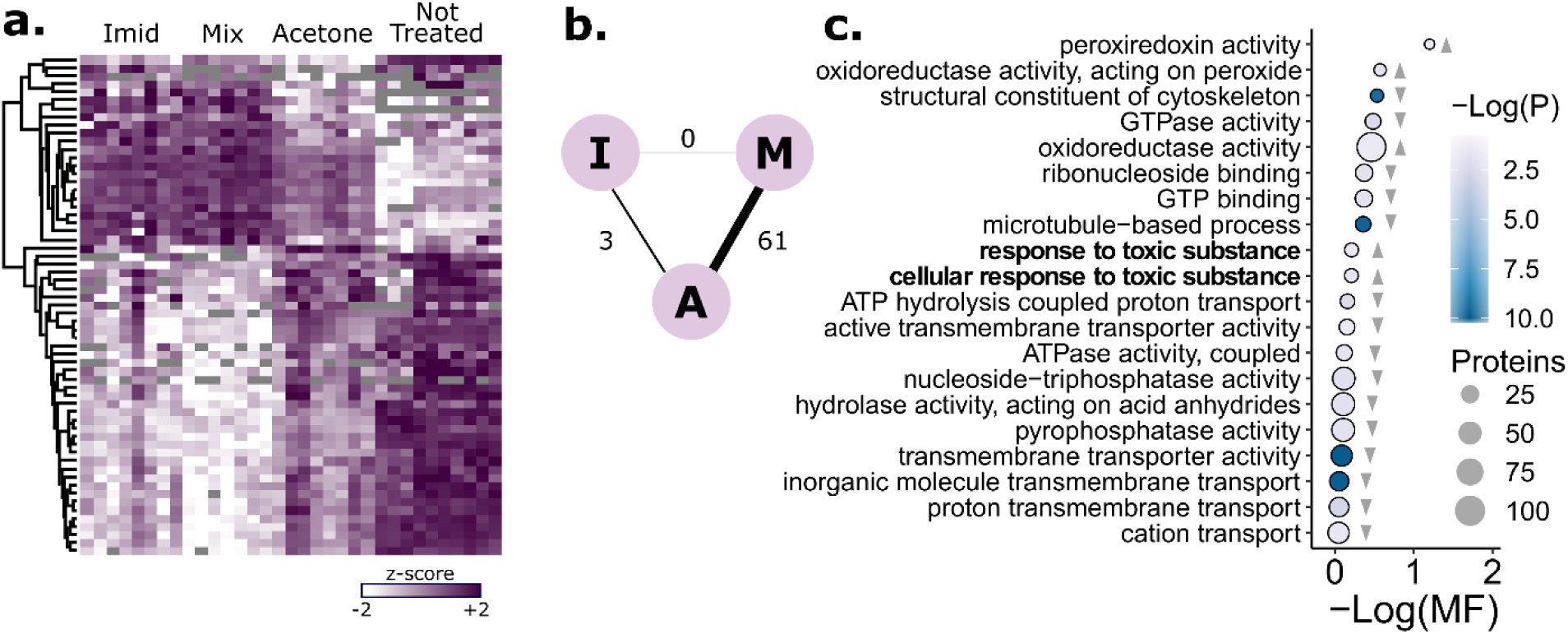
Impact of direct contact pesticide exposure on queen spermathecal protein expression. **A)** Protein expression changes induced by pesticide exposure and acetone (only proteins significantly different between the pesticide and acetone treatments, surviving 10% FDR threshold by the Benjamini-Hochberg method, are shown). The pesticide treatment was a low-dose mixture of compounds (acaricides, fungicides, herbicides, and pesticides) found in the wax of honey bee colonies, blended in field-realistic proportions. See **Table 1** for the relative components. The total dose each queen received was equivalent to a hazard quotient of 502 (imidacloprid) or 511 (mixture). Each column represents the spermatheca from one queen. **B)** Schematic of differentially expressed proteins between groups (A = acetone, I = imidacloprid, M = mixture). **C)** GO term enrichment analysis. Grey triangles indicate the direction of expression driving the enrichment (pesticide mixture relative to acetone). Only GO terms passing a 5% false discovery cut-off are shown (Benjamini-Hochberg method). MF stands for multifunctionality score (higher values are less multifunctional). Circle size is proportional to the number of proteins belonging to the GO term.

Of 2,408 quantified proteins, 61 were differentially expressed between the pesticide cocktail-treated and acetone-treated queens (10% FDR, Benjamini-Hochberg correction). Although imidacloprid has a unique mode of action relative to the compounds included in our pesticide treatment mixture, queens exposed to the imidacloprid treatment yielded highly similar protein expression patterns when compared to cocktail-treated queens, and no proteins were differentially expressed between the two treatment groups. Despite showing the same trends as the cocktail treatments, only three proteins were differentially expressed between the imidacloprid and acetone groups, apparently owing to higher variability in the imidacloprid samples (**Figure 3b**). A single protein, XP_625100.1 (a glyoxylase), was differentially expressed in both imidacloprid- and cocktail-treated queens relative to acetone queens. As expected, the GO term ‘response to toxic substance,’ driven by upregulated proteins, was significantly enriched among others (**Figure 3c**). The top two most significantly upregulated proteins in the pesticide cocktail-treated queens were XP_026296889.1 (catalase, a well-known peroxidase) and XP_392368.1 (cytochrome c oxidase subunit 5a, the final complex in the electron transport chain).

### Expression patterns in failed and healthy queens

To observe proteomic shifts in queens that failed in the field, we solicited samples of healthy (n = 45), imported (n = 18), and failed (n = 60) queens from beekeepers in British Columbia, Canada. Failed queens were defined as any queen whose reproductive capacity was not satisfactory to the beekeeper (*e.g*., symptoms include poor laying pattern, poor colony build-up, poor colony strength, and opportunistic diseases). We found a general upregulation of 139 proteins in failed queens (5% FDR, Benjamini-Hochberg correction), and again the GO term ‘response to toxic substance’ was significantly enriched (**Figure 4a and b**). Interestingly, the candidate markers, two for heat-shock and two for pesticide exposure, were significantly upregulated in failed queens relative to controls at a global scale, whereas the cold-shock markers were not (**Figure 4c;** see **Table 2** for summary statistics).

**Table 2.** Summary statistics for candidate stress biomarkers

| Accession | Stress | Comparison | Test statistic | P | Adjusted P |
| --- | --- | --- | --- | --- | --- |
| XP_001120006.2 | Heat | Heat-shock:Control | 2.89 | 0.00044 | 0.013 |
| | | Failed:Healthy | 4.51 | $1.72 \times 10^{-5}$ | 0.0021 |
| XP_395659.1 | Heat | Heat-shock:Control | 3.22 | 0.0015 | 0.022 |
| | | Failed:Healthy | 5.26 | $8.50 \times 10^{-7}$ | 0.00024 |
| XP_026296889.1 | Pesticide | Pest:Acetone | 3.9 | 0.00183 | 0.085 |
| | | Failed:Healthy | 6.47 | $2.59 \times 10^{-9}$ | $3.27 \times 10^{-6}$ |
| XP_392368.1 | Pesticide | Pest:Acetone | 4.54 | 0.000554 | 0.045 |
|  |  | Failed:Healthy | 3.13 | 0.00234 | 0.045 |
| XP_026296654.1 | Cold | Cold-shock:Control | 5.54 | 0.00025 | 0.042 |
|  |  | Failed:Healthy | 2.29 | 0.024 | 0.17 |
| XP_395122.1 | Cold | Cold-shock:Control | 5.63 | 0.00032 | 0.038 |
|  |  | Failed:Healthy | 0.74 | 0.46 | 0.74 |

**Figure 4.**
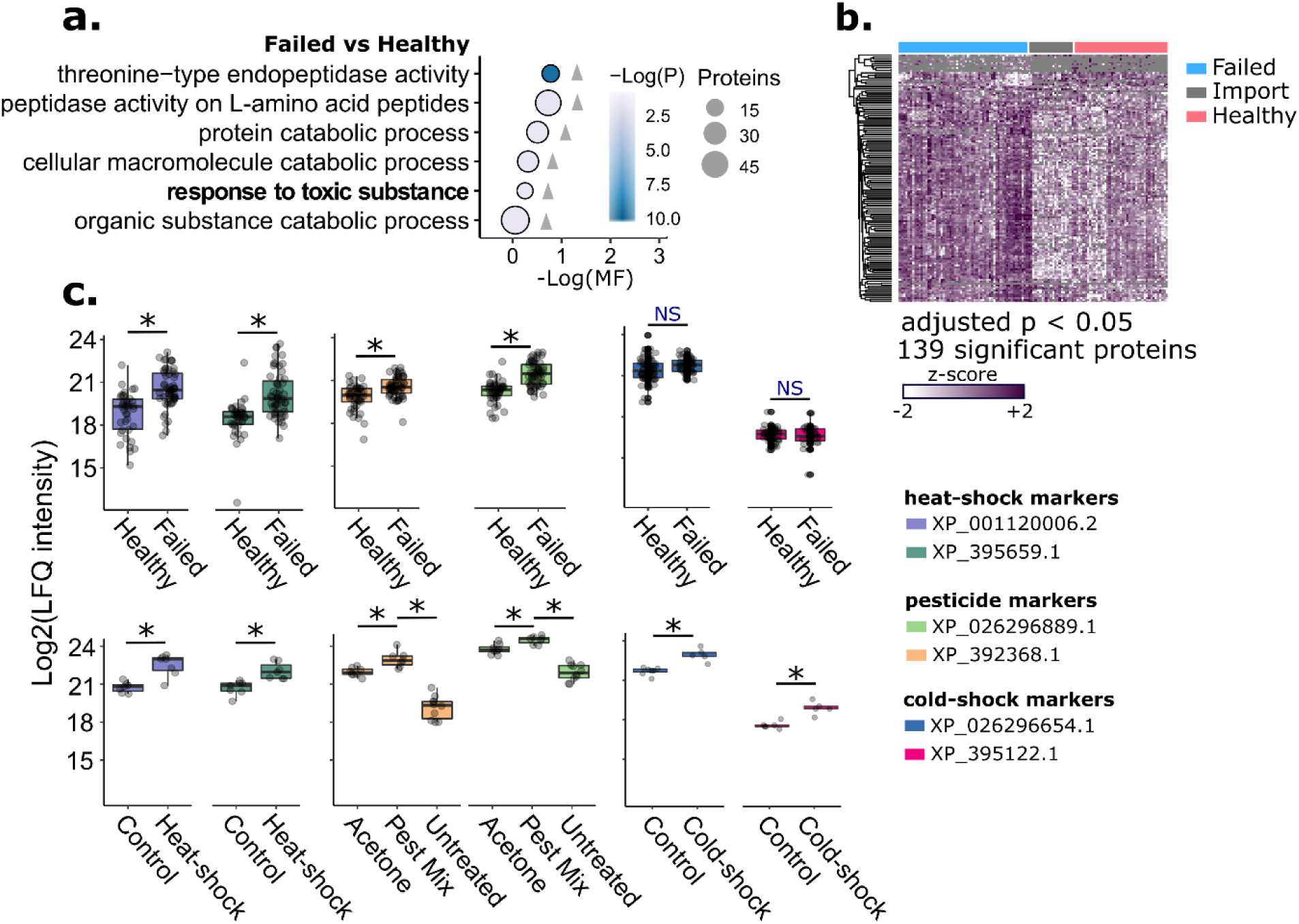
Expression patterns of candidate biomarkers in healthy and failed queens sampled from beekeepers. **A)** GO term enrichment analysis. Grey triangles indicate the direction of expression driving the enrichment (failed relative to healthy). Only GO terms passing a 5% false discovery cut-off are shown (Benjamini-Hochberg method). MF stands for multifunctionality score (higher values are less multifunctional). Circle size is proportional to the number of proteins belonging to the GO term. **B)** Protein expression patterns of failed queens (N = 60), imported queens from Hawaii (N = 9) and California (N = 9), and healthy queens (N = 45). Only significantly different proteins (5% FDR, Benjamini-Hochberg method) are shown. **C)** The top two most significantly upregulated proteins from each experimental stress condition were chosen as the candidate biomarkers. Plots show their expression patterns in the experimental stress conditions and the healthy vs. failed queens.

**Figure 5.**
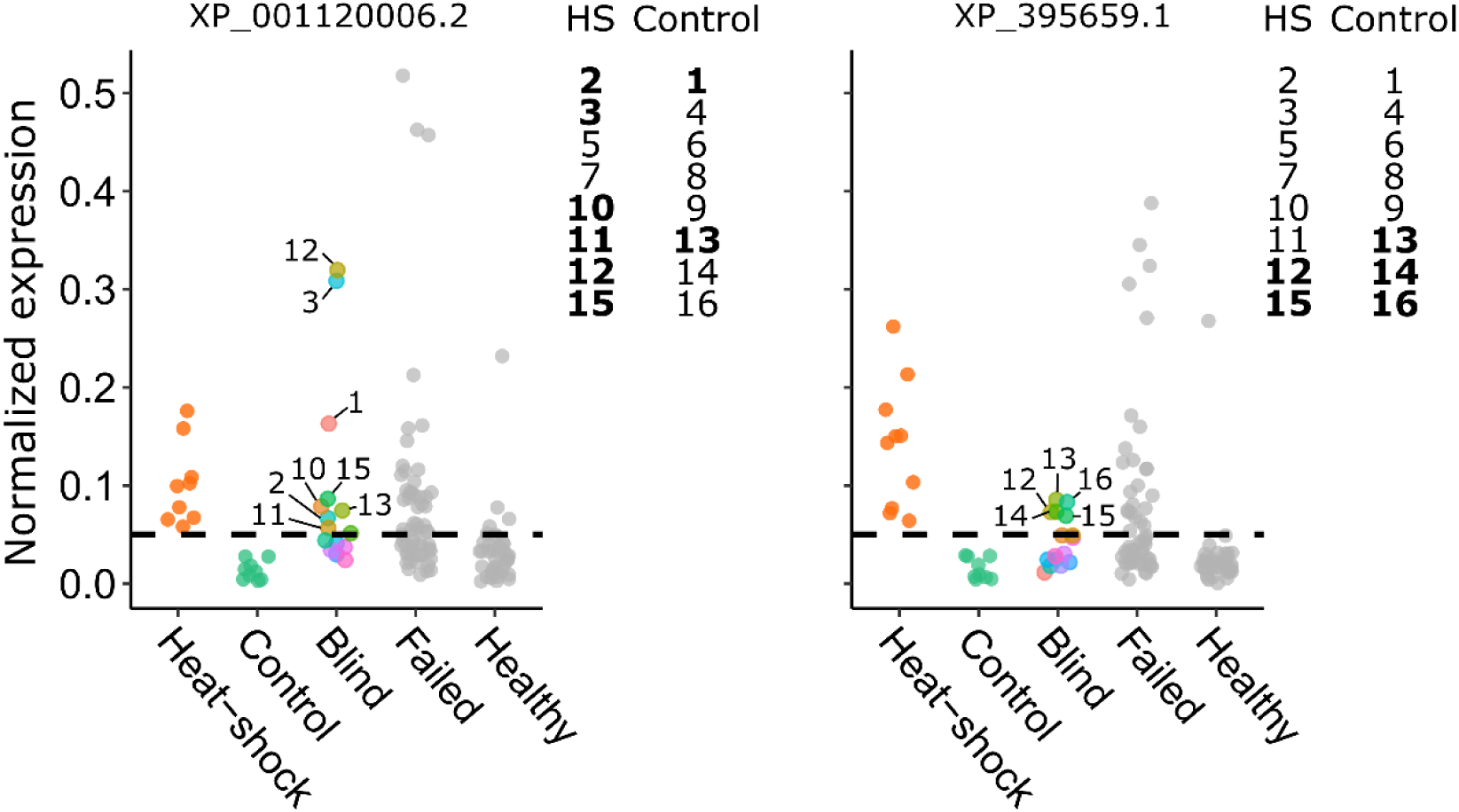
Expression of candidate heat-shock biomarkers in a blind heat-shock trial. Protein expression (normalized LFQ intensity) was normalized to the protein with the lowest variance across all the spermatheca samples (XP_623495.1, V-type proton ATPase catalytic subunit). The heat-shock and control data were previously published. The blind queens were randomized into experimental groups (heat-shock (HS) and control) by a third party and groupings were only revealed after the data were plotted. Queens in the HS group were exposed to 40 °C for two hours, whereas queens in the control group were held at 27 °C. The prediction accuracy of candidate marker XP_001120006.2 outperformed the candidate marker XP_395659.1.

### Blind heat-shock trial

To determine if we could use the heat-shock markers, XP_001120006.2 and XP_395659.1, to correctly identify heat-shocked queens despite no knowledge of their treatment group, we performed a blind heat-shock trial. Setting the expression elevation threshold to 0.05 (only heat-shocked queens showed expression above this threshold in the previous unblind trial), eight of sixteen queens displayed elevated XP_001120006.2 (six of which were correctly assigned), whereas five queens displayed elevated XP_395659.1 (only two of which were correctly assigned). Therefore, XP_001120006.2 is a better biomarker, with 75% true positive rate, whereas XP_395659.1 performed poorly, with only 25% true positive rate.

## Discussion

Here we describe progress made on identifying biomarkers for different forms of queen stress, which we ultimately aim to develop as a diagnostic test. We report new candidate markers proposed for cold-stress and pesticide-stress, and further develop those we have already proposed for heat-stress. Through this shot-gun discovery approach, we also gain some insight into how these stressors affect the biology of the spermatheca on a proteome-wide scale. Interestingly, some of these proposed markers are also upregulated in queens that failed passively in the field, indicating that they could have practical utility for the apiculture industry.

Both cold-shock and heat-shock caused general down-regulation of proteins involved in small molecule metabolism (*e.g*., carboxylic acids and amino acids) and oxidoreductase activity, whereas for cold-shock, proteins involved in lipid metabolic processes increased. It is unclear why fatty acid metabolism is favored by cold-shock, as is the role of the two cold-shock biomarkers we proposed: leucine-rich repeat transmembrane neuronal protein 4-like (or LRRT-4, XP_026296654.1) and probable tRNA N6-adenosine threonylcarbamoyltransferase (XP_395122.1). LRRTs are best known for their roles in regulation and development of excitatory and inhibitory synapses [24]. However, our sample preparation method should not enable extraction of neuronal proteins nor transmembrane proteins (it is the soluble spermathecal fluid proteome), so this function is unlikely here. Indeed, inspecting the sequence using the Basic Local Alignment Search Tool (BLAST) shows that while the protein does have an LRR domain, most of the protein’s sequence is devoted to an E3 ubiquitin-protein ligase domain (which tags proteins with ubiquitin, targeting them for degradation). Therefore, we think it is more likely that XP_026296654.1 is involved in the downregulation of oxidoreductases and small molecule catabolic enzymes that we observe with cold-shock. It is not clear what the relationship is between the threonylcarbamoyltransferase enzyme (which, as the name suggests, adds a threonylcarbamoyl group to specific tRNAs) and cold-shock. But, overall, these results are consistent with temperature stress generally causing metabolic dysregulation in the spermatheca, which may contribute to how heat-and cold-shock leads to sperm death.

The imidacloprid pesticide treatment was originally included only as a positive control, since contact with such a large dose inside a colony is highly unlikely (and therefore, we did not aim to find queen biomarkers for exposure to this pesticide). We expected the imidacloprid treatment to have the greatest effect on protein expression; however, the expression patterns of the imidacloprid-treated queens were remarkably similar to the cocktail-treated queens. This is despite the cocktail not containing any imidacloprid (indeed, it does not contain any neonicotinoid insecticides), and accounting for the effect of the carrier (acetone) by making statistical comparisons only to acetone-only controls (and not untreated queens). Our data, therefore, show a generalized detoxification response that was unselective when compared between a compound with a distinctive mode of action (imidacloprid) and a nine-component pesticide mixture. It could be that other tissues (*e.g*., the fat body and hemolymph) are the major sites of detoxification from contact exposures, and the changes observed in the spermathecal fluid proteome are mainly secondary effects. Indeed, while ‘response to toxic substance’ was a significantly enriched GO term, other GO terms were far more significant (‘structural constituent of cytoskeleton,’ ‘microtubule-based process,’ and ‘transmembrane transporter activity’) were highly significantly enriched via protein downregulation.

Catalase, which was upregulated in pesticide-treated queens, is well known for its role in mitigating peroxide-induced oxidative stress, and its upregulation has been previously linked to pesticide exposure in other insects (*e.g. Harmonia axyridis, Sogatella furcifera, Diadegma semiclausum*) [25-27]. This is likely because pesticide detoxification often leads to oxidative stress, which must be counteracted to avoid DNA damage and cell death [28-30]. The observation that not only catalase, but also other enzymes contributing to the ‘response to toxic substance’ GO term were upregulated in both the pesticide-treated and the failed queens suggests that failed queens may suffer from increased exposure to environmental toxins, possibly through chronic lifetime contact exposure to pesticides in wax. Although the exact causes of failure for these queens are ultimately unknown, these observations are consistent with previous work linking queen failure, reduced brood production, and loss of sperm viability to pesticide exposure (indeed, these failed queens also had significantly lower sperm viability compared to healthy queens, as we previously reported) [5, 7, 21, 31].

Although catalase was also upregulated in failed queens as well as experimentally pesticide-stressed queens, this does not necessarily mean that the failed queens were pesticide stressed. Similarly, although the candidate heat-shock markers were upregulated in failed queens, at this stage we cannot say definitively if this was related to heat. This is because the failed queens also tended to be older than the healthy queens, and because of this sampling bias, we cannot confidently distinguish between proteins changing as a result of age, and those changing with health status. However, even if catalase expression is mainly influenced by age, that would also be an interesting result. In other species (including other insects), catalase expression is linked to fecundity and is thought to help protect the developing oocyte from oxidative damage [32]. Stored sperm require similar protection from oxidative stress (indeed, honey bees have evolved other strategies, such as a highly anaerobic spermathecal environment, to help achieve this) [33-35]. But catalase expression typically declines with age, along with fecundity [32], which is not the trend we observed here. If honey bee queens have instead evolved to upregulate catalase with age (with or without added pesticide stress), this could help explain how they maintain high fecundity throughout their long lives, through improved protection of oocytes and sperm cells from oxidative damage. Indeed, catalase expression appears to be tightly linked to sperm maintenance, as the enzyme becomes significantly upregulated in the spermatheca during the transition from virgin to mated queen [7, 34, 35]. More experiments will be necessary to determine whether catalase expression correlates with age independently of queen health status.

In the blind heat-shock trial, we were surprised that the candidate biomarkers (the small heat-shock proteins (HSPs) XP_001120006.2 and XP_39659.1) were not more useful for classifying queens into their respective treatment groups. We expected that queens with both proteins significantly upregulated would be, most confidently, the ones that were heat-shocked; however, of the three queens with both proteins upregulated, only two were actually heat-shocked. Therefore, this method has lower true positive rate (25% compared to 75%) than using XP_001120006.2 alone. Although a 75% true positive rate is better than chance, it is still not satisfactory for a diagnostic test. For the test to be practically useful, we aim to explore different methods of data acquisition and normalization to minimize noise, as well as determine the biological limits of marker utility (*e.g*., how long after heat-shock the marker remains upregulated).

Important biomarker attributes remain to be determined, especially for the cold-stress and pesticide-stress candidates, which are yet lacking dose-response tests. For the markers to be practically useful, the minimum stress threshold leading to marker activation needs to be determined (similar to how we determined that heat-stress for >1 h is necessary to activate the sHSP markers, here) through dose-response experiments. Such experiments would also help identify if, for example, in extreme cases, new, more determinant biomarkers are activated. Likewise, their practical utility will also be influenced by the time window between which the markers are activated and when they decline back to basal levels (if ever), especially considering that the main test subjects will be failed queens – a phenotype which can take months post-stress to observe. For factors such as pesticide stress, where queens are likely chronically exposed, such a decline in marker expression may not even occur. However, it is not known exactly how much exposure queens experience from chronic contact with contaminated wax.

## Conclusion

Heat-shock, cold-shock, and pesticide exposure induce unique proteomic stress signatures in honey bee queen spermathecae. In addition to offering biological insights into how these stressors alter the spermathecal environment, and providing candidate proteins to investigate as potential causal factors for stress-associated reductions in sperm viability, these signatures may also be useful as biomarkers for specific causes of queen failure. Some of these protein expression patterns are also apparent in queens that failed in the field, but due to sampling bias, we cannot determine the extent to which these expression patterns are linked to failure, and the extent to which they are linked to age. Our blind heat-shock trial shows that markers with similar stress-associated expression patterns can have markedly different predictive power, demonstrating that each marker will need to be individually optimized and validated before they can be used as a reliable diagnostic test.

## Methods

### Queen sources

Queens for temperature-stress tests were shipped to Vancouver, Canada, on July 3^rd^, 2019, from a commercial supplier in California. Queens for the pesticide-stress experiments were shipped on April 24^th^, 2019, from a commercial supplier in Chile. Immediately upon arrival, queens and attendants were given a few drops of water and held overnight at 30°C before exposure to their respective stressors. Queens for the survey of failed and healthy queens were obtained from local beekeepers and queen breeders in British Columbia, Canada (Kettle Valley Queens, Nicola Valley Honey, Wild Antho, Campbells Gold Honey, Heather Meadows Honey Farm, Six Legs Good Apiaries, Wildwood Queens, Cariboo Honey, and Worker Bee Honey Company) between the first week of June 5^th^ and July 10^th^, 2019. See **Table S1** for supplementary queen information and descriptions (such as shipment method, mating day, lineage, etc.). Queens were shipped with temperature loggers (WatchDog B1000 series) to verify that there was not adverse exposure during transport. ‘Healthy’ queens were all evaluated to have a good laying pattern by the beekeepers (consistent egg distribution, few missed cells, one egg per cell) and were two months old. ‘Failed’ queens were rated as inferior by beekeepers on the basis of symptoms such as poor colony population build-up, drone-laying, and sparse brood patterns [36]. Some were failing due to diseases such as chalkbrood or had EFB-like symptoms, and it was not clear if this was an effect of queen quality or some other factor. We previously verified that failed queens had significantly lower sperm viability than healthy queens [6]. Failed queens tended to be older than healthy queens, and ages of some were unknown. Imported queens were obtained from Hawaii (n = 9) and California (n = 9), shipped by international air, arriving on June 14^th^, 2019.

### Stressing queens

For the time-course heat-shock experiment, queens were held in an incubator set to 40 °C and 40% RH for 0.25, 0.5, 1, or 2 h (n = 7 each). Another group of queens (n = 6) were held at 40 °C for 2 h at 80% RH to test for a humidity effect. For the cold-shock experiment, queens (n = 5) were held at 4 °C for 2 h, then 30 °C for 2 d. Control queens (n = 7) were not exposed to heat, but were held in the same 30 °C incubator for two days with the other queens.

Queens for the pesticide-stress experiments were first anesthetized with carbon dioxide, then 2 µl of acetone containing either nothing, 20 ppb imidacloprid (Chem Service Inc, West Chester, PA), 97.9% purity, 20 ppb produced via serial dilution in acetone), or a mix of nine different pesticides listed in **Table 1**, yielding a cumulative HQ of 502, was dispensed onto the thorax of each queen (n = 8, 8, and 7, respectively). All compounds in the pesticide mixture were purchased as pure technical material (≥95% purity) from Sigma Aldrich (St. Louis, MO) or Chem Service Inc and were serially diluted in acetone in order to achieve the respective concentrations listed in **Table 1**. An additional group of n = 10 queens were untreated. Queens were then held at 30 °C for 2 d prior to dissecting and freezing their spermathecae.

### Blind heat-shock trial

Queens for the blind heat-shock experiment were removed from active hives near Kutztown, PA and were 1-6 months of age and actively laying. The colonies were rated as being in good health with no visual signs of disease and good brood pattern. Queens were placed in queen cages with attendants and food and transported via automobile to NC State University within two days of being removed from colonies. Queens were placed inside the aluminum block well of a dry heat bath with cover and held for two hours at 40 °C. Queens were then held in an incubator for an additional 48 h at 30 °C and then frozen at −80 °C and shipped on dry ice to British Columbia, Canada.

### Proteomics sample preparation

Protein was extracted from spermathecae by lysing the organ in 100 µl Buffer D (17 mM D-glucose, 54 mM KCl, 25 mM NaHCO_3_, 83 mM Na_3_C_6_H_5_O_7_). Sperm cells and tissue debris were spun down by centrifuging tubes at 2,000 g for five minutes, then pipetting off the top 80 µl of supernatant. The supernatant was diluted 1:1 with distilled water, then proteins were precipitated by adding four volumes of ice-cold 100% acetone and incubating at −20°C overnight. Precipitated proteins were spun down (10,000 g for 10 min, 4°C) and the supernatant was removed. The protein pellets were washed twice with 500 µl ice-cold 80% acetone, dried at room temperature for 15 min, then solubilized in urea digestion buffer (6 M urea, 2 M thiourea, 100 mM Tris, pH 8.0) and processed for proteomics analysis as previously described [7]. Briefly, protein samples were reduced (1 µg dithiothreitol), alkylated (5 µg iodoacetamide, dark), and digested (1 µg Lys-C, 4 h, then 1 µg Trypsin, overnight) at room temperature. Peptides were acidified, then desalted using in-house packed C18 STAGE tips, quantified using a nanodrop (A280 nm), and 1 µg of peptides were analyzed in random order by liquid chromatography-tandem mass spectrometry (LC-MSMS; Thermo Easy-nLC 1000 coupled to a Bruker Impact II quadrupole time-of-flight mass spectrometer) [37].

### Mass spectrometry data processing

Raw mass spectrometry data files were searched with MaxQuant (v 1.6.1.0) [38] using default parameters except label-free quantification was enabled, match between runs was enabled, and the LFQ min ratio count was set to 1. Complete search parameters can be found within the mqpar.xml file within the Proteome Exchange repository. The *Apis mellifera* reference proteome database was downloaded from NCBI on November 18^th^, 2019 (HAv3.1) and all honey bee virus sequences were added to the fasta file. The final search database is also available in the data repository. All data associated with this manuscript (raw data files and search result files) are available on ProteomeXchange (www.proteomexchange.org; accessions: *upload in progress*).

### Differential expression analysis

Differential expression analysis for the experimentally stressed queens was performed using Perseus (v 1.5.6.0) as previously described [7]. Briefly, data were log2 transformed, then reverse hits, proteins only identified by site, and contaminants were removed, along with any protein identified in fewer than six samples. Samples were compared in pairwise combinations (heat-shock to control, cold-shock to control, and pesticide-stressed to acetone) using t-tests with a Benjamini-Hochberg multiple hypothesis testing correction to 5% FDR (for heat-shock and cold-shock experiments) or 10% FDR (for the pesticide-stress experiment). For the pesticide experiments, we used acetone-treated queens as the negative control because we were interested in genes that were influenced by the pesticides specifically, over and above the effect of the solvent. However, we still evaluated expression in untreated queens in order to confirm that proteins differentially expressed between acetone and pesticide queens were also differentially expressed between pesticide queens and untreated queens, and in the same direction (representing the more natural condition). For this experiment, we relaxed the FDR because at 5% FDR there were not two proteins significantly up-regulated compared to acetone samples which were also identified in the healthy vs. failed queen survey.

The queen survey data is part of a larger project (yet unpublished) evaluating relationships between spermathecal protein expression and other queen phenotypes; therefore, these data were analyzed with a more sophisticated linear model with multiple fixed continuous and discrete covariates. To do this, we used the limma package in R [39]. The results of this analysis were also corrected to 5% FDR by the Benjamini-Hochberg method.

### GO term enrichment analysis

Gene ontology (GO) terms were retrieved for protein groups using BLAST2GO (v 4.0) and enrichment tests were conducted using the gene score resampling method within ErmineJ [23]. This method operates independently of a ‘hit list’ of significantly differentially expressed proteins, and is therefore not sensitive to sliding FDR thresholds of up-stream differential expression analyses. Rather, it reads in raw p values and calculates if proteins with low p values are more likely to share GO terms than expected by chance. More documentation about the gene score resampling method can be found on the software developer’s website [15]. For each experimental condition, we performed enrichment analyses on upregulated and downregulated proteins separately. The enrichment FDR was also adjusted to 5% using the Benjamini-Hochberg method.

## List of abbreviations

BLAST: (basic local alignment search tool)
DMPF: (2,4-Dimethylphenyl-N’-methyl-formamidine (degradate of Amitraz))
FDR: (false discovery rate)
GO: (gene ontology)
HQ: (hazard quotient)
HSP: (heat-shock protein)
LC-MSMS: (liquid chromatography tandem mass spectrometry)
LFQ: (label-free quantification)
LRRT: protein (leucine-rich repeat transmembrane protein)
MF: (multifunctionality)
RH: (relative humidity)

## Declarations

### Ethics approval and consent to participate

As a non-cephalopod invertebrate species, honey bees are not subject to animal ethics committee approval at the University of British Columbia.

### Consent for publication

Not applicable.

### Availability of data and materials

The datasets and databases generated and/or analysed during the current study are available in the Pride ProteomeXchange mass spectrometry data repository, www.proteomexchange.org. Accessions: *upload in progress*).

### Authors’ contributions

AM wrote the manuscript, designed the experiments, prepared proteomics samples, and analyzed the proteomics data. AC coordinated and received queens for the queen survey and dissected queens for the stress tests. JM produced the pesticide cocktail, helped design the pesticide stress experiment, and assisted with data analysis and interpretation. JSP conducted the blind heat-shock trial. Grants to AM, LJF, JSP, and DRT funded the research.

## Acknowledgements

We thank Kettle Valley Queens, Nicola Valley Honey, Wild Antho, Campbells Gold Honey, Heather Meadows Honey Farm, Six Legs Good Apiaries, Wildwood Queens, Cariboo Honey, and Worker Bee Honey Company for donating failed and healthy queens for this research. We also thank Robyn Underwood for supplying the queens used for the blind heat-shock test, and Heather Higo, Marta Guarna, and Liz Huxter for organizing the BC queen survey.

## Footnotes

Not applicable.

